# Genomic resistance in historical clinical isolates increased in frequency and mobility after the age of antibiotics

**DOI:** 10.1101/2025.01.16.633422

**Authors:** Arya Kaul, Célia Souque, Mische Holland, Michael Baym

## Abstract

Antibiotic resistance is frequently observed shortly after the clinical introduction of an antibiotic. Whether and how frequently that resistance occurred before the introduction is harder to determine, as isolates could not have been tested for resistance before an antibiotic was discovered. Historical collections, like the British National Collection of Type Cultures (NCTC), stretching back to 1885, provide a window into this history. Here we match 1,817 sequenced high-quality genomes from the NCTC collection to their respective year of isolation to study resistance genes before and concurrent with the age of antibiotics. Concordant with previous work, we find resistance genes in both pathogens and environmental samples before the age of antibiotics. While generally rare before the introduction of an antibiotic, we find an associated increase in frequency with antibiotic introduction. Finally, we observe a trend of resistance elements becoming both increasingly mobile and nested within multiple mobile elements as time goes on. More broadly, our findings suggest that likely-functional antibiotic resistance genes were circulating in clinically relevant isolates before the age of antibiotics, but human usage is associated with increasing both their overall prevalence and mobility.

**DATA SUMMARY:** Genome assemblies downloaded and analyzed are in Supplementary Table 1, and computational tools used are found in the Methods. **The authors confirm all supporting data, code and protocols have been provided within the article or through supplementary data files.**

**Impact statement:** Historical collections of microbial isolates enable researchers to both investigate the past and identify interesting trends over time. In this study, we queried over 1,800 isolate genomes in one such collection for genomic variation linked to antibiotic resistance. We show that numerous isolates cultured before the introduction of a given antibiotic contain genomic variation linked to antibiotic resistance; however, this phenomenon remained relatively rare. We demonstrate a strong association between the year a given antibiotic was clinically introduced and a rise in prevalence of genomic resistance to that antibiotic. Finally, we show that while mobile elements are common throughout the isolates and timeframe analyzed, genomic resistance has become increasingly mobile as time has gone on. This study shows that as expected, the clinical introduction of a given antibiotic is correlated with an increase in resistance to that antibiotic but also was linked with increased mobility of genes and alleles conferring resistance. However, we note that the effect of deposition bias in the collection cannot be excluded. Our work also indicates that numerous microbial pangenomes of pathogens naturally contained genomic resistance to a given antibiotic even before anthropogenic use of that antibiotic. Taken together, we demonstrate that although human use may affect the prevalence and mobility of genomic resistance in clinical isolates; for most antibiotics, genomic resistance existed within the pangenomes of sampled pathogens prior to clinical introduction. Quantifying and understanding the impact of antibiotic introduction in the past helps us understand how the introduction of novel antibiotics can impact bacteria; allowing better reaction to novel resistant infections as they arise.

## INTRODUCTION

Since the earliest days of modern antibiotics, their use has been threatened by resistance (1). Bacteria acquire antibiotic resistance, the ability to survive a higher dose than can be safely given to a patient, both by mutating their genomes and by acquisition of new genes. The use of antibiotics then selects for resistant mutants, allowing them to preferentially replicate and spread. Indeed, for every antibiotic introduced, resistance soon follows.

To understand the evolutionary history of clinical resistance, we not only need to understand the selective pressures on resistant mutants, but also what the frequency and prevalence of resistance alleles was before the widespread use of antibiotics. This presents a problem: how can we know the rate of resistance to an antibiotic before it was even discovered? Contemporaneous with early antibiotic discovery, freeze-dried collections of bacterial isolates began to be collected and preserved to this day. In 1920, the United Kingdom’s National Collection of Type Cultures (NCTC) was founded, preserving both clinical and environmental isolates for long-term storage (2). Prior studies have leveraged these collections to gain insights from the phylogeography of individual strains to retrospective diagnoses of ancient disease outbreaks (3–5). More recently, large scale sequencing efforts have produced many publicly available, high-quality genomic assemblies from the NCTC, which we use here to study resistance alleles around the advent of antibiotics (6).

While most explanations for antibiotic resistance attribute the problem to human use of antibiotics, suggesting that consumption and production of these drugs have created selective pressure driving bacteria to develop and retain resistance, recent studies have identified antibiotic resistance in microbes isolated before the widespread use of antibiotics.

Most explanations for antibiotic resistance attribute the problem to human use of antibiotics, suggesting consumption and production of these drugs have created selective pressure that drives bacteria to develop and retain resistance (1). Recent work studying microbes isolated before the widespread use of antibiotics has found antibiotic-resistance even in clinical isolates. The first NCTC isolate accessioned, NCTC1, a *Shigella flexneri* isolated in 1915, is resistant to penicillin and erythromycin, well before the discovery of either drug (7,8). Phylogenetic reconstruction has predicted methicillin-resistant *Staphylococcus aureus* first emerged in the mid-1940s, about 14 years before the introduction of methicillin in the clinic (9). In the environment, genomic studies of microbial isolates from permafrost have identified functional antibiotic resistance genes well over 30,000 years old (10,11). While this natural occurrence is unsurprising, given that microbial antibiotic production far predates human use, the frequency, type, and context of resistance mutations in human-associated bacteria before the use of antibiotics remains unknown (10).

Here, we sought to identify and quantify trends in resistance genes contemporaneous with antibiotic introduction in genomes from the NCTC. We note that the present study, by virtue of being solely computational, only considers the presence of genomic resistance determinants. Carriage of a known resistance gene or allele does not necessarily imply resistance to that antibiotic. For example, if a resistance gene is unexpressed, the microbe may still be antibiotic sensitive. However, carriage of a resistance element enables a larger pool of possible mutations which induce phenotypic resistance (12).

We measured and characterized the genomic resistance that existed before the age of antibiotics to better understand how the clinical introduction of a given antibiotic correlates with differences in the frequency and features of resistance. Through the construction and analysis of a time-matched database of microbial genomes we show that human introduction of an antibiotic is associated with changing both the prevalence and mobility of genomic resistance elements within isolates deposited in the NCTC.

## METHODS

### Creation of a time matched database of microbial genomes

To generate a historical database of microbial genomes, we turned to the NCTC administered by the UK Health Security Agency (UKHSA). We chose the NCTC due it its status as the oldest strain collection in the world; and as a result, is one of the most diverse collection of isolates cultured before the age of antibiotics (13). We note that although the database has a predominantly clinical focus, there are also numerous environmental isolates deposited.

To generate the database, we systematically analyzed the web metadata associated with each isolate from the NCTC webserver, as no complete database of isolates is publicly available (https://culturecollections.org.uk). Upon downloading each isolate’s metadata, we aggregated the metadata and then performed a regex search for a putative year of isolation (Supplemental Table 1). Due to the NCTC’s long history, standardized data entry fields and data logging practices are not common throughout the database. If a year of isolation is recorded, it can appear in several different fields. From the regex search, we take all prospective years of isolation and rank order the priority of each year based on a heuristic determined from direct communication with the NCTC. Although a ‘collection date’ field exists for assemblies deposited in GenBank we found it to be either inaccurate: out of 2,958 assemblies in the NCTC3000 project, 1,845 have non-empty collection dates, and only 590 have a single year listed in the collection_date entry. We also found several strains with demonstrably incorrect collection_date entries. For example, NCTC11317 is listed as being isolated in 1969, while the GenBank entry for it lists the collection_date as 2015. In another instance, NCTC13760 is only listed as being “pre-1989” but has a collection_date entry of 1949. For this reason, we neglected to use the collection_date entry and relied on the NCTC metadata associated with each to determine a year of isolation.

For all strains for which a year of isolation was determined, we searched for complete genome assemblies within the European Nucleotide Archive (ENA), Refseq, and Genbank. We found that if a given accession had a genome assembly in either Refseq or Genbank, then there was a copy also deposited within the ENA. For consistency, we only moved forward with assemblies downloaded from the ENA. This output was then merged and analyzed (Figure 1). Out of the 1,919 genomes in our database, 1,752 are from the NCTC3000 collection (91%). The NCTC3000 was generated from a collaboration between the NCTC, Pacific Biosciences, and the Wellcome Sanger Institute which sequenced and assembled around 3,000 NCTC isolates using long-read technology (6).

**Figure 1:**
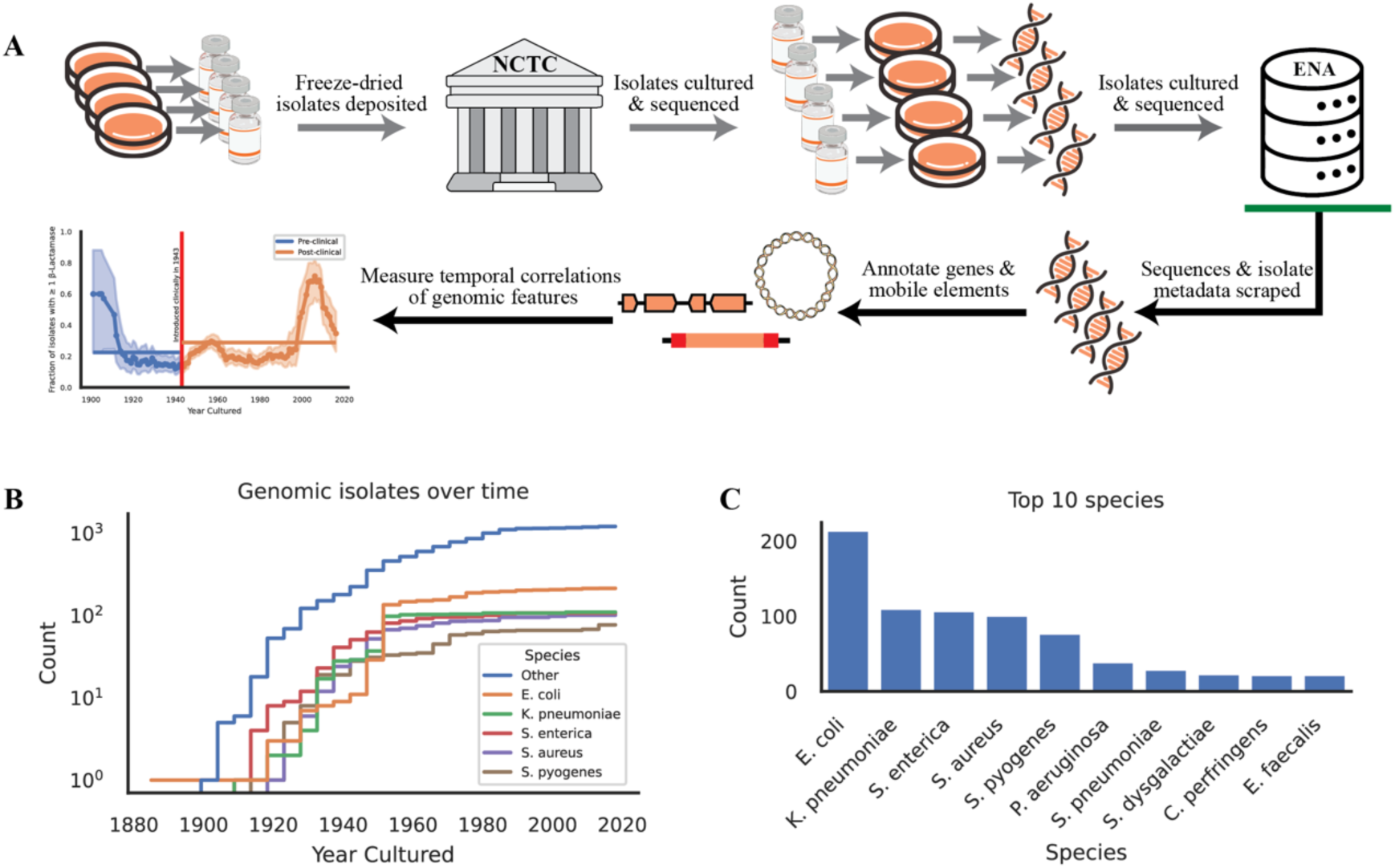
Overview of the collated database. (A) Schematic of the workflow, arrows in gray are prior work not performed in the paper while arrows after the green bar indicate novel work done in this paper. (B) Cumulative count of the genomic isolates captured over time colored with the top 5 species represented in the database. (C) Raw counts of the top 10 species in the database.

To evaluate the quality of our genomic dataset, we calculated three metrics: CheckM2 completeness and contamination scores, as well as N50 (14). We defined a genome as low-quality if it had a completeness score below 90%, a contamination score above 5%, or an N50 value less than 5 kb. After applying these quality control criteria, we filtered out 102 genomes from our initial dataset of 1,919 and were left with a final set of 1,817 high-quality genomes suitable for downstream analysis.

### Identification of genomic resistance elements

Here we use the term “genomic resistance elements” to mean loci on a genome (either SNPs or complete genes) which have been associated with resistance to antibiotics. While the presence of these genomic resistance elements is not a direct predictor of resistance, their presence does correlate with phenotypic resistance (15). To identify genomic resistance elements within each isolate, we took two orthogonal approaches. First, we used the Resistance Genome Identifier and version 3.2.2 of the Comprehensive Antibiotic Resistance Database (CARD) with the parameters (--input_type contig --exclude_nudge) (16). CARD is an extensive database of curated antimicrobial resistance gene sequences and resistance-conferring mutations and can provide a complete view of genomic resistance elements present within a given genome. Additionally, we used AMRFinderPlus on isolates from one of the following taxa using the --organism flag (*Klebsiella pneumoniae*, *Escherichia coli*, *Salmonella enterica*, *Acinetobacter baumannii*, *Enterococcus faecium*, *Staphylococcus aureus*, *Streptococcus pyogenes*, *Streptococcus pneumoniae*, *Enterobacter cloacae*, and *Pseudomonas aeruginosa*) (17). AMRFinderPlus similarly performs alignment searches in a large database of known resistance-associated alleles. Of interest, when combined with the –organism flag, it also incorporates prior knowledge about which resistance alleles are commonly found in a given taxa and filters those results out. As a result, it should be viewed as a refinement tool, allowing for the identification of resistance alleles specific to each taxon. Results from both approaches are described and it is noted when a given analysis uses one set of results over the other.

### Identification of mobility

To classify plasmids present in the assemblies, we used MOBTyper with default options, and every contig analyzed was classified as deriving from the chromosome or a plasmid (18,19). To identify integrons, we used IntegronFinder with default options (20). To identify transposons (TN) and insertion sequences (IS) present in each isolate, we used mobileelementfinder with default options (21). To identify prophages, we used phigaro with default options (22).

For all genomic resistance elements, we classified it with one or more of the following labels. If the element was present on a contig classified as a plasmid by MOBTyper then it was determined to be ‘plasmid associated’. If 95% or more of the element sequence was encompassed by an integron gene cassette, then it was determined to be ‘integron associated’. If the element was flanked by two transposons or two insertion sequence elements within 10 kb of each other and of the same type, or if the element overlapped with a TN or IS, we classified it as ‘TN/IS associated’. If the element was encompassed inside a prophage or was within 1 kb of a prophage we classified it as ‘prophage associated’. If the element met none of these requirements, then it was classified as ‘Immobile’.

We package this pipeline into a package named ‘CallMeMobile’. This pipeline is available for download and installation on Github (https://github.com/aryakaul/callmemobile).

### Null distribution for genomic resistance prevalence

The year in which a given antibiotic was clinically introduced was determined from Hutchings, et al. 2019 (23). To determine the significance of the observed difference between the fraction of isolates containing a genomic resistance element against an antibiotic before and after its clinical introduction, we generated a null distribution using two shuffling methods. In the first method, all years of isolation were collated, and each isolate was randomly assigned a year of isolation from the pool of possible values without replacement. This unstructured shuffling allowed for isolates from different years to be reassigned to any other year; for example, some isolates from 1958 could be assigned to 2009 while others could be assigned to 1930. To account for possible structure in the year of isolation, we employed a second method: structured shuffling. In this approach, all isolates from a given year were randomly assigned to the same new year. For example, all isolates cultured in 1958 would be randomly assigned to the same year; for example, 1930.

We performed both shuffling methods 10,000 times. A Gaussian distribution was fitted to the created null distribution, and a p-value was determined from where the observed difference fell. The unstructured shuffling always yielded lower p-values due to the uneven density of samples in the NCTC. As such, we only report the more stringent p-values derived from the structured shuffling approach, which appears to better account for these sampling biases.

### Genomic Annotation, Data Processing and Visualizations

Genomes were uniformly annotated with Bakta using version 5 of their database and default options (24). DNA Features Viewer was used to visualize resulting GFFs (25). Dataframes were processed with Pandas and all plots were created with Seaborn (26–28).

### Sampling Bias

As with any analysis of a historical database, sampling bias is a significant consideration. This analysis using the NCTC is no exception. First, the NCTC is based in the United Kingdom and administered by the UK Health Security Agency. As a result, most isolates deposited are either from the UK or Europe and most isolates are clinically relevant. Thus, the isolates included in our study are not necessarily representative of the global bacterial population present during those time periods.

In addition, the availability and preservation of isolates from different time periods is not uniform, with some periods being more highly sampled than others (Figure 1B). This temporal bias could impact our ability to accurately assess the prevalence and characteristics of antibiotic resistance alleles throughout history. In addition, the isolates deposited are reflective of researcher’s priorities at given points in time. For example, there is a noticeable spike in the fraction of isolates with genomic resistance elements deposited after the year 2000; this likely coincides with increased interest in clinical resistance and its markers.

Though we strove to mitigate these biases through random sampling and a focus on species with large sample sizes, we note that the complete elimination of sampling bias is impossible.

## RESULTS

### Creation of an accessible NCTC genomic database with a year of isolation

At the time of download, 5,665 isolates were present in the NCTC. Of these, 1,919 had both a predicted year of isolation from the metadata and publicly available assembled genomes. These 1,919 isolates represent 605 species with the majority (651 isolates) being either *Escherichia coli* (228), *Klebsiella pneumoniae* (122), *Salmonella enterica* (119), *Staphylococcus aureus* (105), or *Streptococcus pyogenes* (77).

We present an interactive Jupyter notebook in the project Github repository to make the database easily accessible for other researchers (https://github.com/baymlab/MicroTrawler). Both the complete data of the 5,665 isolates scraped from the NCTC, the 1,919 genomes scraped, and the 1,817 high quality genomes used for the rest of this paper’s analysis can be found in Supplementary Table 1.

### Genomic resistance elements were present in isolates cultured before the clinical introduction of a given antibiotic

We first measured the prevalence of resistance to a given antibiotic before the clinical introduction of that antibiotic. To focus our analysis on the most informative genomic resistance elements likely to lead to a phenotypic change in resistance profile, we employed AMRFinderPlus and focused on a subset of the species in our database. These included the 274 isolates belonging to one of the ESKAPE pathogens (*Enterococcus faecium* (6), *Staphylococcus aureus* (101), *Klebsiella pneumoniae* (110), *Acinetobacter baumannii* (9), *Pseudomonas aeruginosa* (39), and *Enterobacter* sp. (9)), the 212 isolates from *Escherichia coli*, the 106 isolates of *Salmonella enterica*, the 77 isolates of *Streptococcus pyogenes*, and the 29 isolates of *Streptococcus pneumoniae* bringing the total to 698 isolates. Each of the 698 assemblies and their species information was input to AMRFinderPlus to query for resistance elements not commonly found within these taxa (see Methods). To investigate if these isolates harbored genomic resistance elements before the clinical introduction of an antibiotic class, we computed the fraction of isolates before anthropogenic introduction of a given antibiotic that contain alleles associated with resistance to that antibiotic.

We find that although uncommon, it is possible to detect resistance elements in the genomes of clinically relevant isolates cultured before the clinical introduction of a given antibiotic (Figure 2B, Box 1). Moreover, our analysis reveals differential rates of resistance for different antibiotics and microbial species. When we re-run this analysis using CARD and do not remove resistance elements known to be commonplace, we observe widespread and ubiquitous presence of these elements across various antibiotics (Supplementary Figure 2). This suggests that certain resistance genes have been intrinsic to the genomic repertoire of certain taxa for some time, existing prior to human usage of antibiotics.

**Figure 2:**
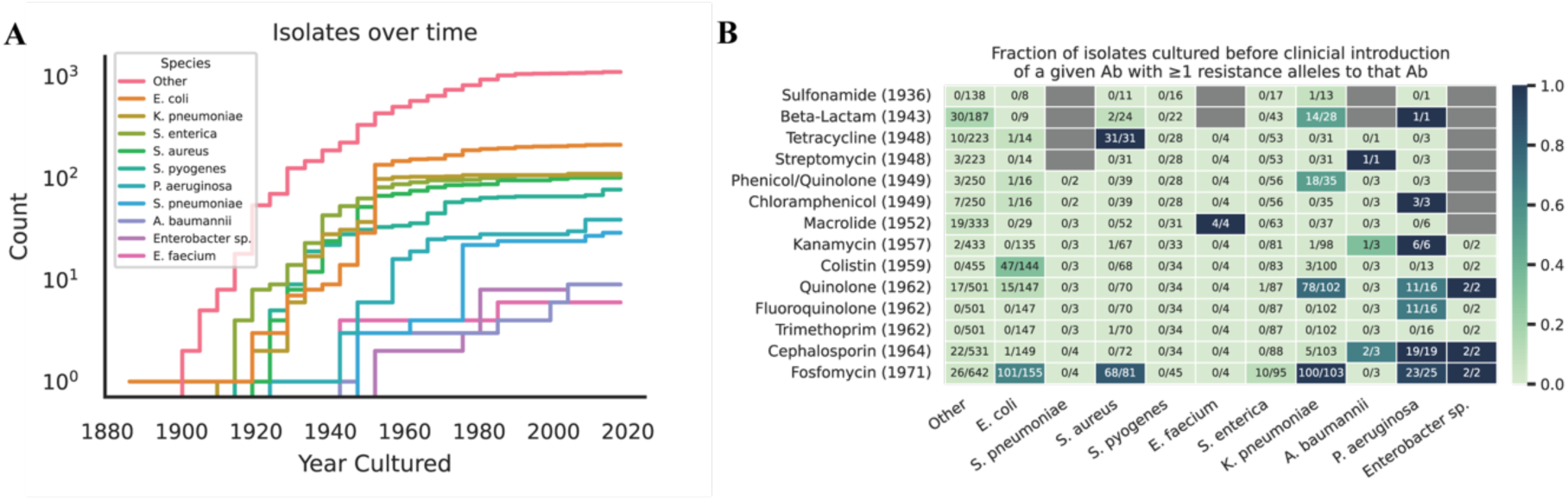
Resistance alleles were present but uncommon across species and drug classes before the introduction of antibiotics. (A) A cumulative histogram of each of the genomic isolates over time in our dataset analyzed with AMRFinderPlus. (B) Heatmap representing the fraction of isolates harboring resistance alleles before the introduction of a given antibiotic. Denominator in each cell is the number of isolates of a given species cultured before the clinical introduction of the antibiotics in the corresponding row (represented by the (Year) next to the antibiotic). The numerator is the number of isolates cultured before clinical introduction of the antibiotic containing resistance alleles to that antibiotic.

To better understand these isolates, we selected a variety of pre-antibiotic era isolates with genomic resistance and analyzed their isolation source and the genomic context of the resistance elements (Box 1, Supplementary Figure 3). Overall, we show that while rare, historical isolates deposited in the context of both clinical and environmental settings contain genomic resistance elements to a given antibiotic, even before clinical usage of that antibiotic.

### Genomic resistance elements against most antibiotics are significantly more prevalent after clinical introduction of that antibiotic

Though we observe some genomic resistance in isolates before the age of antibiotics, we were interested in determining if the clinical introduction of an antibiotic is significantly associated with an increase in the prevalence of genomic resistance to that antibiotic. We began with beta-lactamases, as their prevalence and diversity is well-associated with rising resistance to beta-lactams in the clinic and beta-lactamases are thought to have minimal pleiotropic influences (29,30).

For each year we had isolates from, we measured the fraction of isolates in that year with one or more beta-lactamases. We then took the mean of the fractions before the clinical introduction of penicillin in 1943 and after and measured the difference between the means, denoted by delta. To measure the significance of delta, we shuffled the year of isolation for each isolate and recomputed the delta as before. This null distribution was fitted to a normal distribution and the observed delta’s significance was computed (Methods). We find that the increased fraction of isolates containing beta-lactamases is significantly correlated with the clinical introduction of penicillin in 1943 (Figure 3A). Subsetting the data to only ESKAPE pathogens, provides a more significant difference (Figure 3B). To determine if these results held to broader classes of antibiotics, we expanded this analysis to include all antibiotics for which we had at least 30 resistant isolates overall.

**Figure 3:**
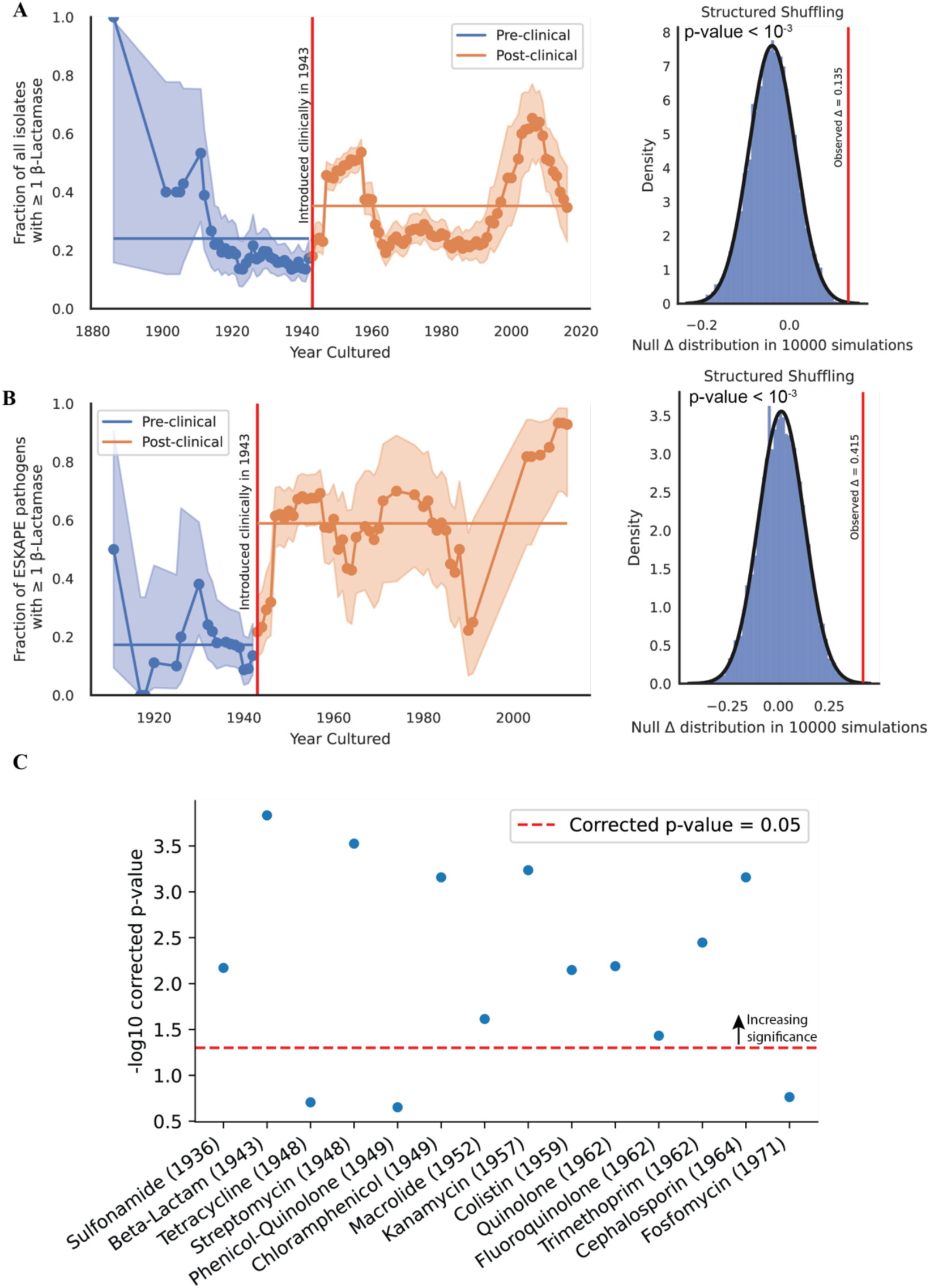
Clinical introduction of antibiotics is significantly associated with an increase in the fraction of isolates containing resistance to that antibiotic across drugs and mechanisms of action. (A) Left panel: Fraction of isolates with ≥ 1 beta-lactamase over time. Error bars represent the beta distribution 95% confidence estimate given the number of isolates in that year. Right panel: the significance of the observed difference in the mean fraction of isolates with ≥ 1 beta-lactamase after 10,000 simulations of randomly shuffling the year of isolation for the data. (B) Same as in (A) but restricted to ESKAPE pathogens. (C) Same analysis performed across all isolates and over all drug classes with 10,000 shuffles. Multiple hypotheses corrected using Benjamini-Hochberg correction.

We find that for most drug classes, the clinical introduction of antibiotics is significantly associated with a rise in resistance alleles present in the NCTC; with the notable exceptions of tetracycline, phenicol/quinolones and fosfomycin (Figure 3C, Supplementary Figure 4).

Pre-1948 tetracycline resistance is driven by the chromosomally encoded efflux pump, tet(38), in *S. aureus* (31). Phenicol/quinolone resistance before 1949 is largely a result of the oqxAB efflux pumps in *K. pneumoniae* (32). Though both genomic elements are thought to only induce low- to medium-resistance to these drug classes, they still constitute clinically relevant genotypes that provide evidence that the pangenome of these isolates often contained genomic resistance elements even before the selective pressure induced by human introduction of antibiotics.

In contrast, fosfomycin resistance pre-dating its clinical introduction in the early 1970s exhibits a broader range of mechanisms. Fosfomycin’s mechanism of action involves inhibiting the enzyme murA, which catalyzes an early step in cell wall synthesis. In some instances, point mutations in glpT and uhpT were observed that decrease the uptake of fosfomycin into the bacterial cell (33). In other instances, murA exhibited mutations known to cause resistance to fosfomycin (34,35) Additionally, the presence of fosA and fosB was noted, which encode enzymes capable of deactivating fosfomycin entirely (36). These genomic determinants of resistance were spread across the species analyzed. We found no association with the change in genomic resistance prevalence when we further stratify the antibiotics by mechanism of action (Supplementary Figure 5).

These results led us to conclude that the human introduction of a given antibiotic is typically followed by the deposition of isolates in the NCTC containing resistance to that antibiotic. We were next interested in investigating how genomic mobility and the capacity for horizontal gene transfer has changed for resistance elements over time.

### Genomic determinants of resistance exhibit increasing mobility over time

To investigate the trends in the association of antibiotic resistance elements with mobile elements over time, we classified each resistance element as either ‘Immobile’ or some combination of ‘Plasmid associated’, ‘Integron associated’, ‘TN/IS associated’, and ‘Prophage associated’ (see Methods).

For those resistance elements that were mobile, we investigated the relative fractions of each of the different mobility types over time. We observe that TN/IS associated resistance elements are the most common, followed by plasmid associated, integron associated, and prophage associated. Importantly, all four seem to show an increase over time, suggesting enhanced potential for horizontal spread of genomic resistance elements among isolates deposited over the course of the NCTC (Figure 4A).

**Figure 4:**
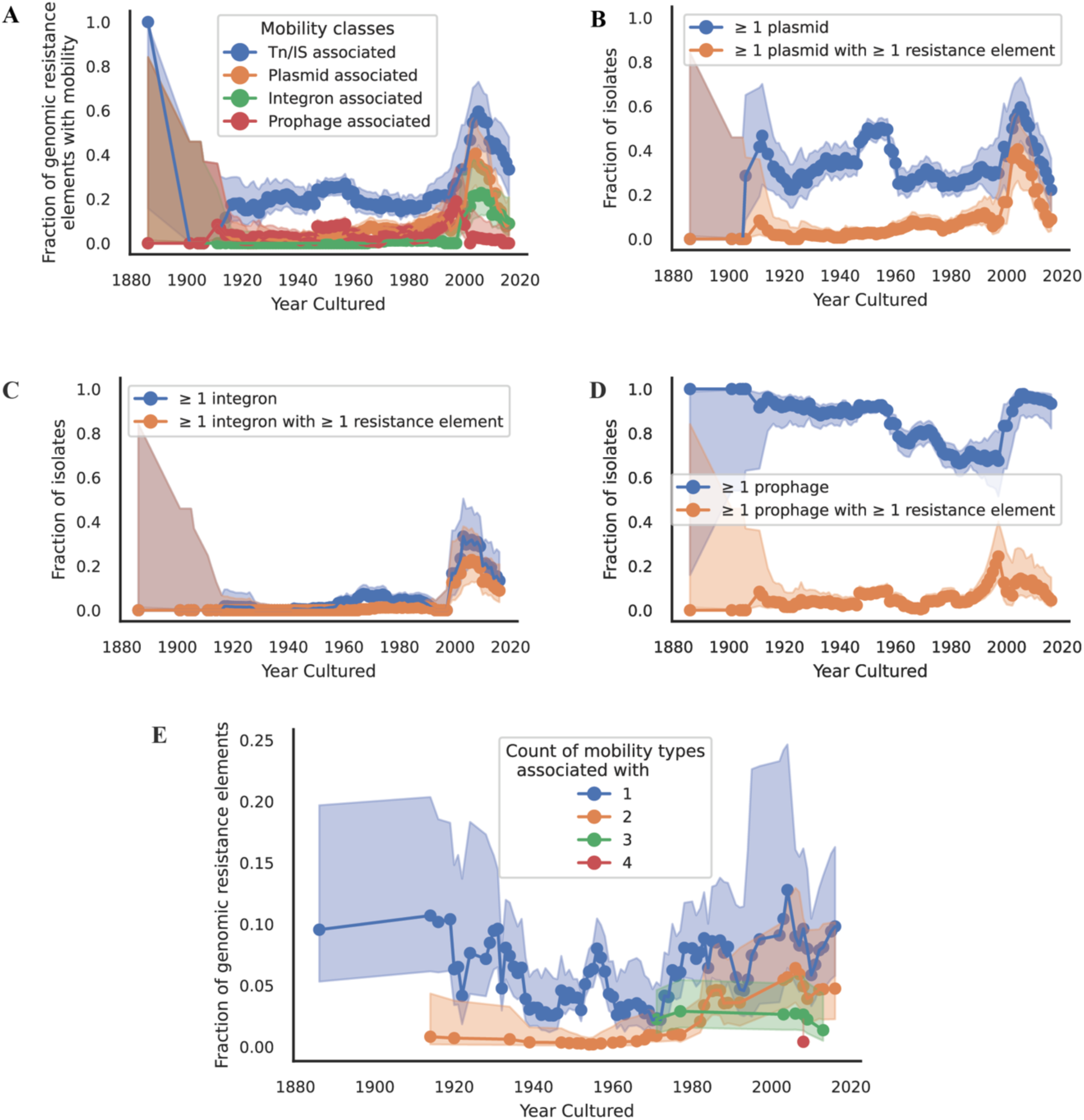
Over time, resistance alleles in the NCTC exhibit increasing mobility. Error bars represent the beta distribution 95% confidence estimate given the number of isolates in that year. (A) The fraction of resistance elements associated with a given mobility type. Note that a given resistance allele might have more than one form of mobility. (B) The fraction of isolates over time that contain one or more plasmids, and the fraction that contain one or more plasmids predicted to harbor one or more resistance elements. (C) Same as (B) but considering integrons. (D) Same as (B) but considering prophages. (E) The fraction of resistance elements with 1 or more independent mobility types associated with it.

To investigate if the cause for this increase in resistance mobility was driven by an increase in mobility overall, we investigated the prevalence of different mobile elements over time, irrespective of them carrying resistance elements. We found that the fraction of isolates containing one or more plasmids is relatively constant over the course of the NCTC (Figure 4B). However, when we then look at the fraction of those plasmids that contain one or more resistance genes we find a steady increase over time, indicating that over the course of the NCTC it becomes more likely to identify isolates with resistance elements carried on plasmids over time (Figure 4B). These findings align with the results from an older work investigating resistance carried by pre-antibiotic era conjugative plasmids, and another more recent study sequencing pre-antibiotic era plasmids and finding an absence of resistance elements (37,38). We performed the same analysis for integrons and prophages. Integrons tended to be rarer in our dataset but do exhibit the same association with resistance elements. We note that since we have fewer integrons in our dataset, these results should be interpreted with caution (Figure 4C). Prophages tended to be abundant across our dataset and only begin to be associated with resistance elements for more recent isolates (Figure 4D).

The previous analysis was done across all genomic resistance elements as identified by CARD. To determine if these same results held for a more focused subset of resistance elements, we again turned to beta-lactamases. Analyzing the mobility of beta-lactamases over time, we find both an overall increase in at least one class of mobility; as well as an increase in plasmid-associated beta-lactamases as time goes on (Supplementary Figure 6).

Finally, we observe a progressive rise in the proportion of resistance genes associated with more than one possible mechanism of mobility (transposons, plasmids, integrons, or prophages) (Figure 4D, Supplementary Figure 6, Supplementary Figure 7). One copy of the dfrA14 gene was found to be associated with all four mobility mechanisms; this variant of dihydrofolate reductase (dfrA) confers resistance to trimethoprim and is commonly linked with Class 1 integrons (39). This dfrA14 was found on NCTC 13440, a *Klebsiella pneumoniae* strain isolated from the Bolzano Regional Hospital in Italy (40). We find it is simultaneously a cassette in a Class 1 integron, within the insertion sequence IS6100, located on a 61kb contig predicted to belong to the conjugative AA552 (IncN) plasmid cluster, and 200bp away from a predicted prophage. Though no other resistance element exhibits all four possible mobility mechanisms, we do note the observed increase in cumulative mobility mechanisms as time goes on. We interpret this increase in mobility as supporting the “nesting doll” pattern of antibiotic resistance, wherein resistance genes become more frequently integrated into multiple mobile genetic elements over time (41).

## DISCUSSION

In Alexander Fleming’s Nobel Prize acceptance speech, he described a model where sub-lethal doses would gradually select for resistance: “Mr. X. has a sore throat. He buys some penicillin and gives himself, not enough to kill the streptococci but enough to educate them to resist penicillin”; in contrast, our study further reinforces that even in the pre-antibiotic era fully resistant pathogens were already likely circulating (42). Rather than de novo resistance emerging in response to antibiotic exposure, we found that antibiotic resistance alleles were already present at low levels in clinically relevant isolates prior to the widespread use of antibiotics. In the metaphor, most likely some subset of streptococci either infecting Mr. X. or in the environment around Mr. X. had already been ‘educated’ to resist the antibiotic. Mr. X.’s use of penicillin merely selects for that subset to proliferate further. This “pre-existing resistance” is consistent with the idea that clinically relevant populations naturally contained genomic resistance elements even before clinical usage of antibiotics (43–45). Though these resistance elements were indeed rare, we note their existence and subsequent rise in frequency after human usage of a given antibiotic.

Furthermore, over time, genomic resistance elements are more likely to be associated with mobile genetic elements. Mobile genetic elements play a critical role in the horizontal transfer of antibiotic resistance genes between bacterial strains (46,47). The increasing mobility of genomic resistance elements over time suggests the accumulation and dissemination of resistance determinants within bacterial populations through horizontal gene transfer mechanisms. This phenomenon has significant implications for the rapid spread of antibiotic resistance, as mobile elements can facilitate the transfer of resistance genes between different bacterial species and genera. In addition, we find evidence for the “nesting doll” theory of antibiotic resistance, where resistance elements become increasingly nested within multiple mobility mechanisms as time has gone on (41,48). This accrual of different mobility mechanisms could be due to homologous recombination between mobile elements and the selection of those mobile elements that carry resistance elements as cargo.

We note two drawbacks to the present study. First, by virtue of being solely computational we cannot make claims about the phenotypic resistance of any of the isolates analyzed. Even though our analysis is relegated to genomic resistance, we argue genomic resistance serves as the raw material for evolution to act upon and is necessarily correlated with phenotypic resistance. Second, all historical analyses of archived isolates are biased by what samples were chosen to be significant enough to add to that archive. We do not believe this bias impacts our first main finding that genomic resistance was present at low levels in clinically relevant isolates even before the introduction of those antibiotics since pre-discovery there was no reason to preferentially deposit resistant isolates. However, we do acknowledge this bias likely skews the further results of changes in resistance prevalence and mobility over time. Though the specific magnitudes and proportions may be influenced by sampling bias, we believe the overall trends observed likely still hold true despite this bias.

More broadly, what we call a resistance gene is itself teleological: while to us their primary role is to resist the antibiotics we use to treat patients, they may have evolved for an entirely different purpose (49). Efflux pumps, for example, confer resistance to a range of natural stressors, and the presence of these genes in bacterial populations prior to the widespread use of antibiotics suggests they were already fulfilling important physiological roles.

The presence of resistance elements in a subsample of the older isolates in the NCTC indicates that they were likely a regular, if infrequent, occurrence in infections before the use of antibiotics. Consistent with our common understanding, the introduction of antibiotics into clinical environments accelerated the prevalence and mobility of these once-rare elements, contributing to the widespread prevalence of resistance observed today.

Our findings add to the growing literature showing that the genetic foundations of antibiotic resistance did not arise solely in response to human antibiotic use (7,10,11). Rather, they were drawn from a long-standing repertoire of resistance alleles already present in the pangenome of clinically relevant isolates. Nevertheless, the introduction of antibiotics into clinical environments does appear to be followed by an acceleration of the prevalence and mobility of these once-rare elements in the NCTC. Recognizing this broader context reinforce that the emergence of clinically relevant antibiotic resistance is not simply a recent phenomenon triggered by modern drug use; rather it is rooted in ancient genomic adaptations that have been amplified by the selective pressures introduced through widespread antibiotic deployment.

## Acknowledgments

We thank the UK Health Security Agency and the National Collection of Type Cultures for providing access to their data and expertise, which guided our research. We thank Fernando Rossine and the entirety of the Baym Lab, whose insights and thoughtful commentary were instrumental in shaping our findings. This work was supported by the NIGMS of the National Institutes of Health (R35GM133700), the Pew Charitable Trusts, and the Alfred P. Sloan Foundation. AK acknowledges support from the NIH NHGRI (T32HG002295). Elements of Figure 1A adapted from CC-BY artwork by the DataBase Center for Life Science, Firefox, Matanbz, Zahra Ibrahem, and Laboratoires Servier.

### Box 1

#### 1.5.1 MdtM gene in a clinically relevant *Salmonella enterica* strain isolated in 1911

NCTC 74 is an isolate of *Salmonella enterica* serotype typhi isolated from a case of food poisoning in 1911 (50). MdtM is a multi-drug transporter efflux pump associated with increased resistance to ciprofloxacin, norfloxacin, levofloxacin, kanamycin, streptomycin, gentamycin, nalidixic acid, chloramphenicol, ethidium bromide, and acriflavine, including fluoroquinolone antibiotics, which are common drugs to treat *Salmonella enterica* typhi infections (51,52).

#### 1.5.2 OqxAB efflux pump in a *Klebsiella pneumoniae* strain isolated in 1920

NCTC 418 is an isolate of Klebsiella pneumoniae deposited within John Hopkins Hospital since at least 1911 (53). The only accompanying metadata associated with it is that it “probably a descendant of the original capsule bacillus of Pfeiffer.” Pfeiffer’s capsule bacillus is now known as *Haemophilus influenzae*; however, comparing k-mer sketching distance of the genome against Refseq genomes shows strong similarity to *Klebsiella pneumoniae* isolates indicating this classification is likely incorrect. NCTC 418 carries the oqxAB operon encoding a small drug efflux pump enabling the bacterium to expel a broad range of toxic compounds. The oqxAB efflux pump has been shown to confer resistance to various antimicrobial agents, including quinolones and biocides (54).

#### 1.5.3 Intact plasmid-borne beta-lactam resistance system in a clinically relevant *Staphylococcus aureus* strain isolated in 1932

NCTC 4136 is an isolate of *Staphylococcus aureus* isolated from a case of food poisoning in 1932 (55). Genomic analysis of its sequence demonstrates the presence of intact BlaR1, BlaZ, and BlaI genes in its genome. In addition, both PlasmidFinder and mob_recon predict that the contig these genes are found on is a plasmid. Annotations on the contig indicate presence of replication proteins and an origin of replication (Supplementary Figure 2C). BlaI serves as the repressor of the operon, while blaR1 senses beta-lactams and blaZ is a beta-lactamase (56).

#### 1.5.4 Mobile sul2 gene in an environmental *Klebsiella pneumoniae* strain isolated in 1933

NCTC 9127 is derived from a study conducting an analysis on the contamination of ice cream in Iowa (57). Later identified as *Klebsiella pneumoniae*, we find 100% sequence homology to the reference sul2 gene deposited in the CARD database. On the bacterial chromosome (contig of 5.3Mb length) we identify sul2, a sulfa-resistant analog for dihydropteroate synthase. In addition, we identify the typical dihydropteroate synthase gene in *Klebsiella pneumoniae*, folP, located upstream of sul2. Consistent with prior literature, we found glmM, a phosphoglucosamine mutase, to be downstream of sul2 (58). We additionally found that both sul2 and the glmM protein sit inside an ICE element known to typically carry both genes, indicating that the horizontal transfer of sulfonamide resistance likely predates the clinical usage of sulfonamide (Supplementary Figure 2D).

#### 1.5.5 Resistant PmrB in a clinically relevant *Escherichia coli* strain isolated in 1947

NCTC 8603 is an isolate of *Escherichia coli* isolated from a case of infantile diarrhea in 1947 that was part of a broader outbreak happening across residential homes and hospitals (59). PmrB is the histidine kinase of the 2-component system PmrAB that regulates the pmrHFIJKLM (or arnBCADTEF) operon (60), and mutations within it have been shown to induce modifications to lipid A that decrease the binding affinity of colistin, a last-line antibiotic (60,61). The E123D mutation noted in NCTC 8603 has not been experimentally confirmed to increase MIC; however, it has been independently observed in colistin-resistant clinical *Escherichia coli* isolates cultured from both swine and humans (62,63).

## Supplementary Figures

**Supplementary Figure 1:**
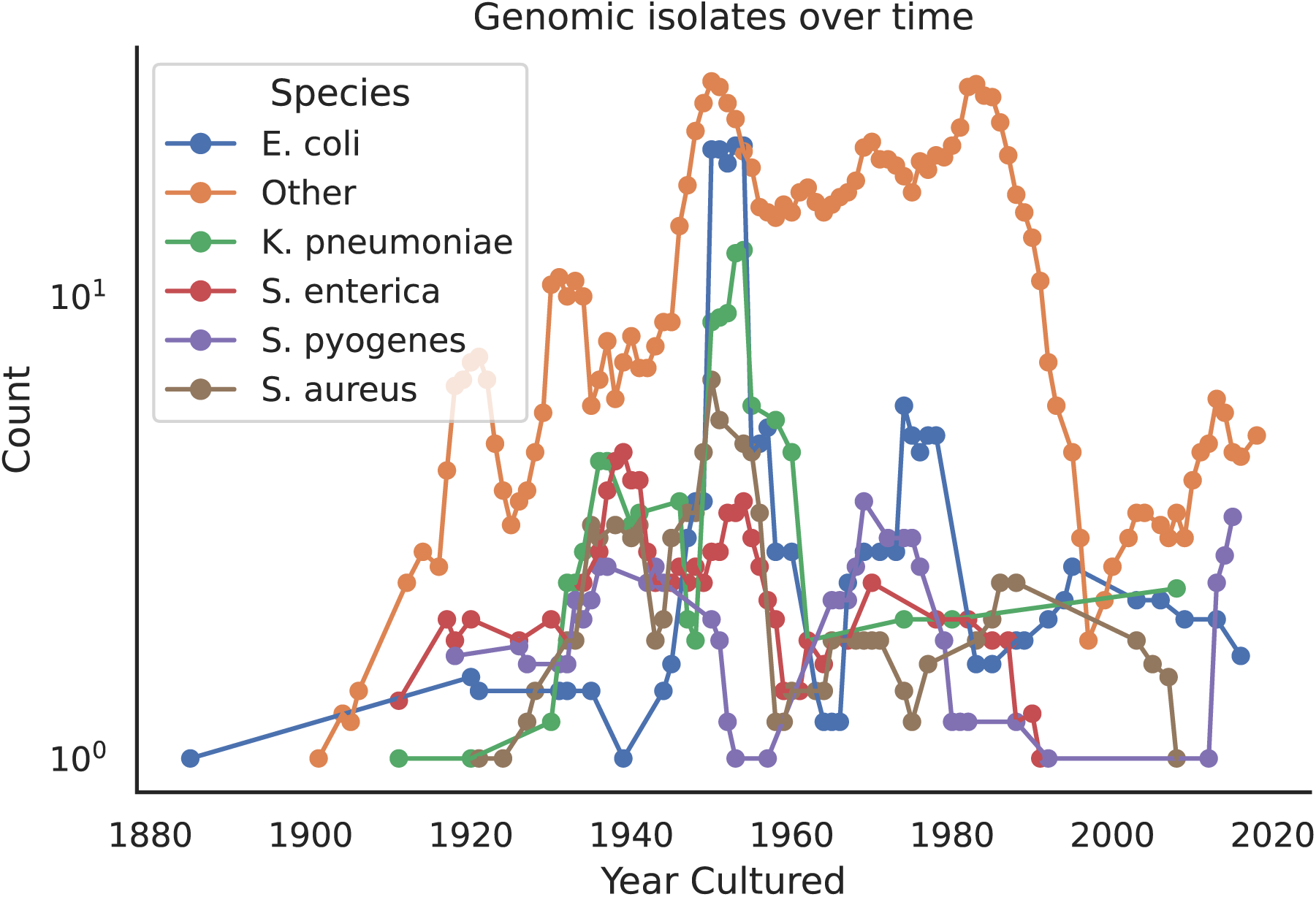
Rolling count of isolates in the dataset over time. Rolling average of number of isolates per specie in the data analyzed. Points represent the average number of isolates isolated within a window 5 years.

**Supplementary Figure 2:**
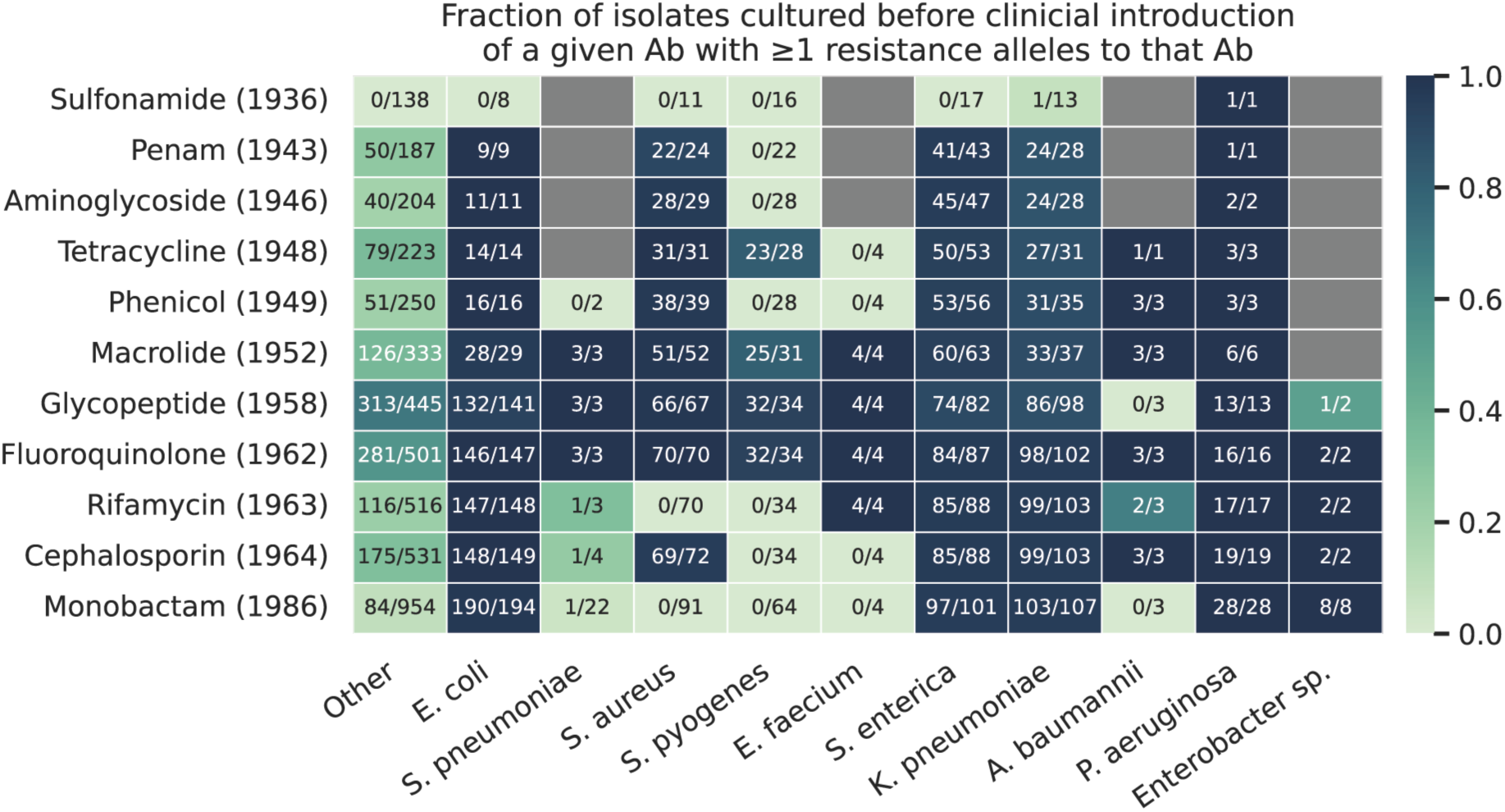
Resistance-associated genomic elements known to exist within specific species’ were also ubiquitous before clinical introduction of given antibiotics. Heatmap representing the fraction of isolates harboring resistance alleles before the introduction of a given antibiotic. Denominator in each cell is the number of isolates of a given species cultured before the clinical introduction of the antibiotics in the corresponding row (represented by the (Year) next to the antibiotic). The numerator is the number of isolates cultured before clinical introduction of the antibiotic containing resistance alleles to that antibiotic. Resistance allele prediction and classification done from CARD without prior species knowledge.

**Supplementary Figure 3:**
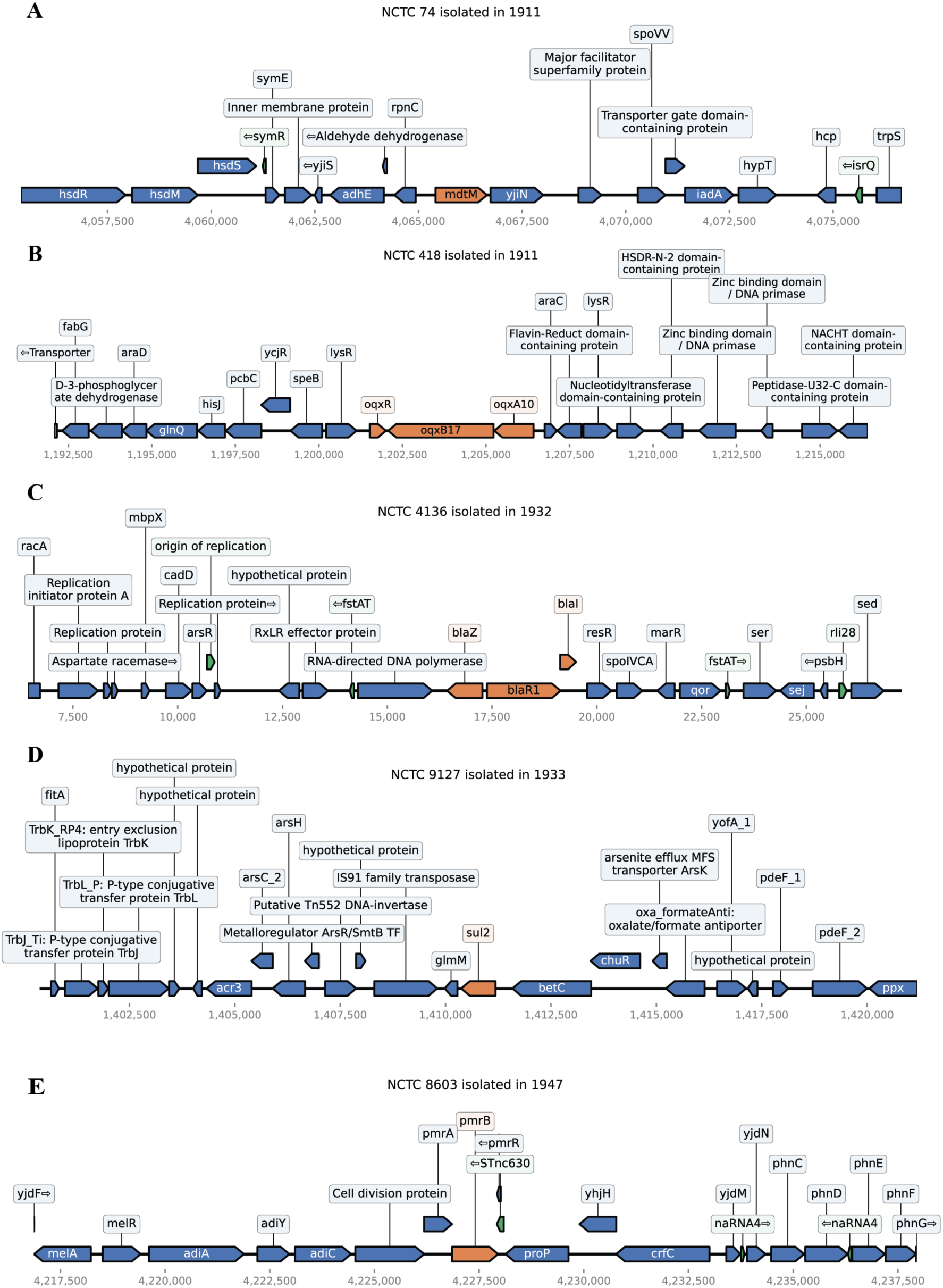
Genomic neighborhood of pre-antibiotic era genomic resistance elements reveals similar genomic structures observed for these resistance alleles. 10kb genomic window surrounding each of the pre-antibiotic resistance elements in the isolates described in Box 1.

**Supplementary Figure 4:**
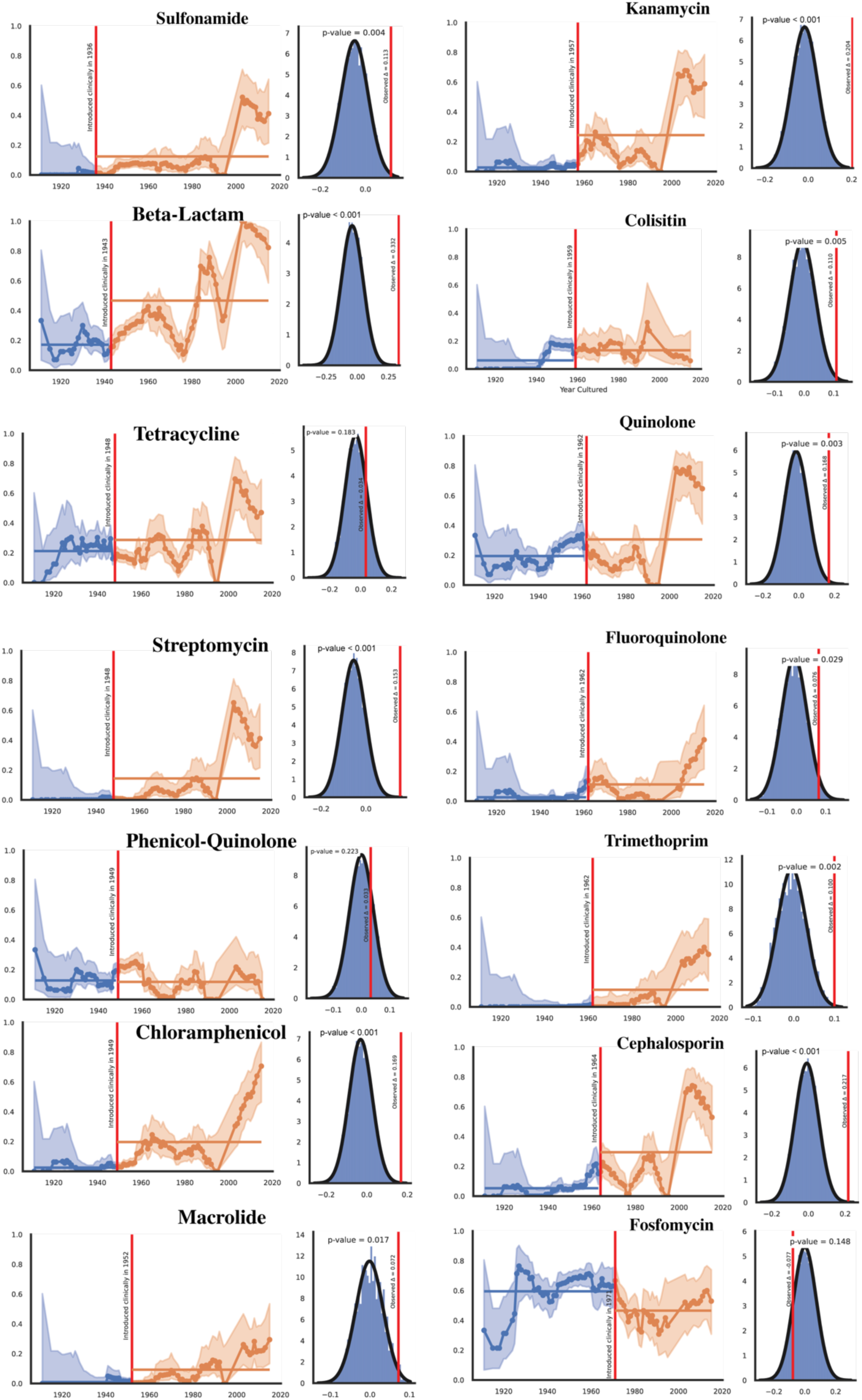
Significant increase in observed resistance frequencies after clinical introduction of an antibiotic across most drug types. See Figure 3 for plot explanation.

**Supplementary Figure 5:**
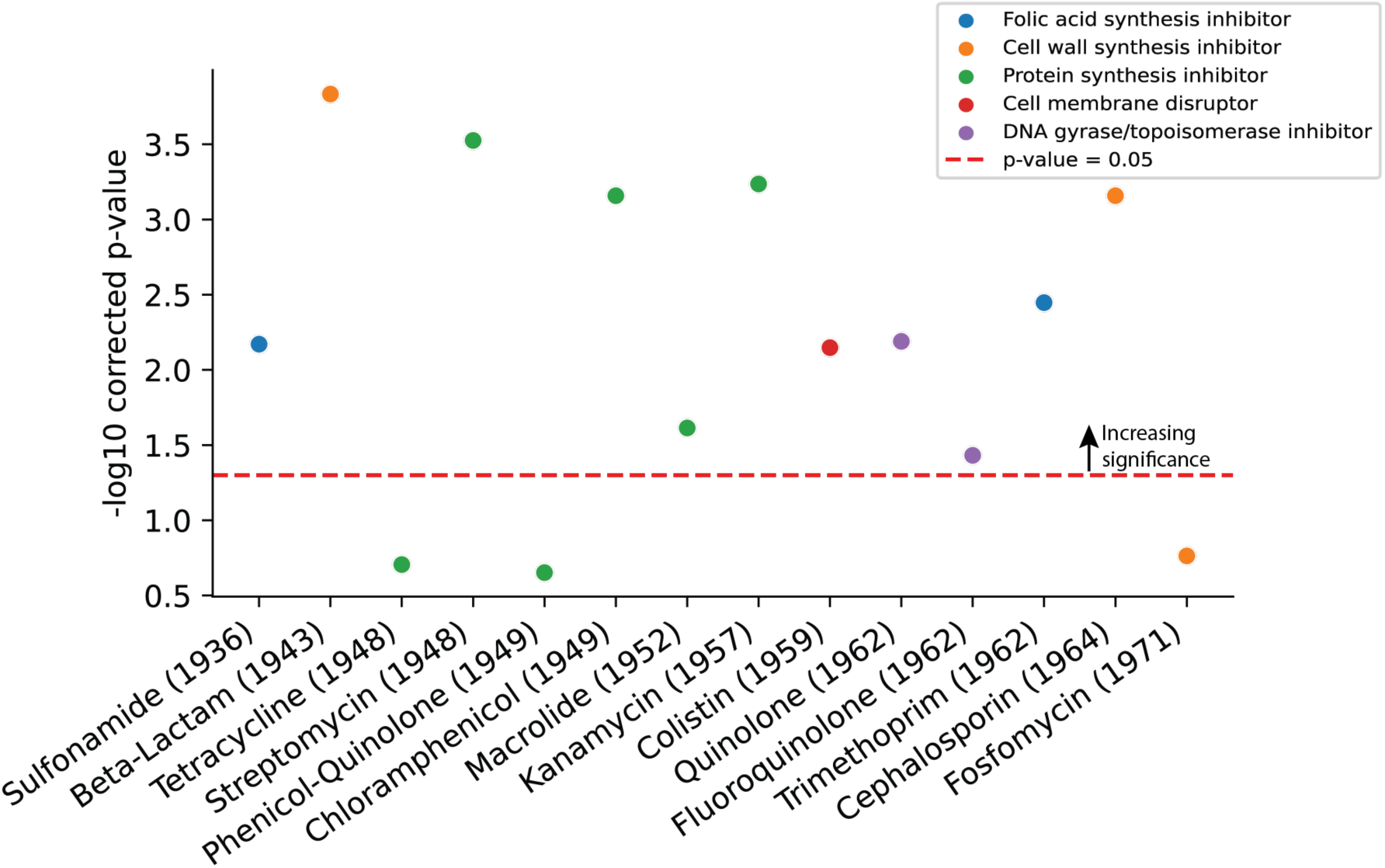
Significant increase in observed resistance rates after clinical introduction are not associated with antibiotic mechanism. Significance of change in resistance prevalence over all isolates and drug classes with 10,000 shuffles. Multiple hypotheses corrected with Benjamini-Hochberg. Individual points colored by antibiotic mechanism.

**Supplementary Figure 6:**
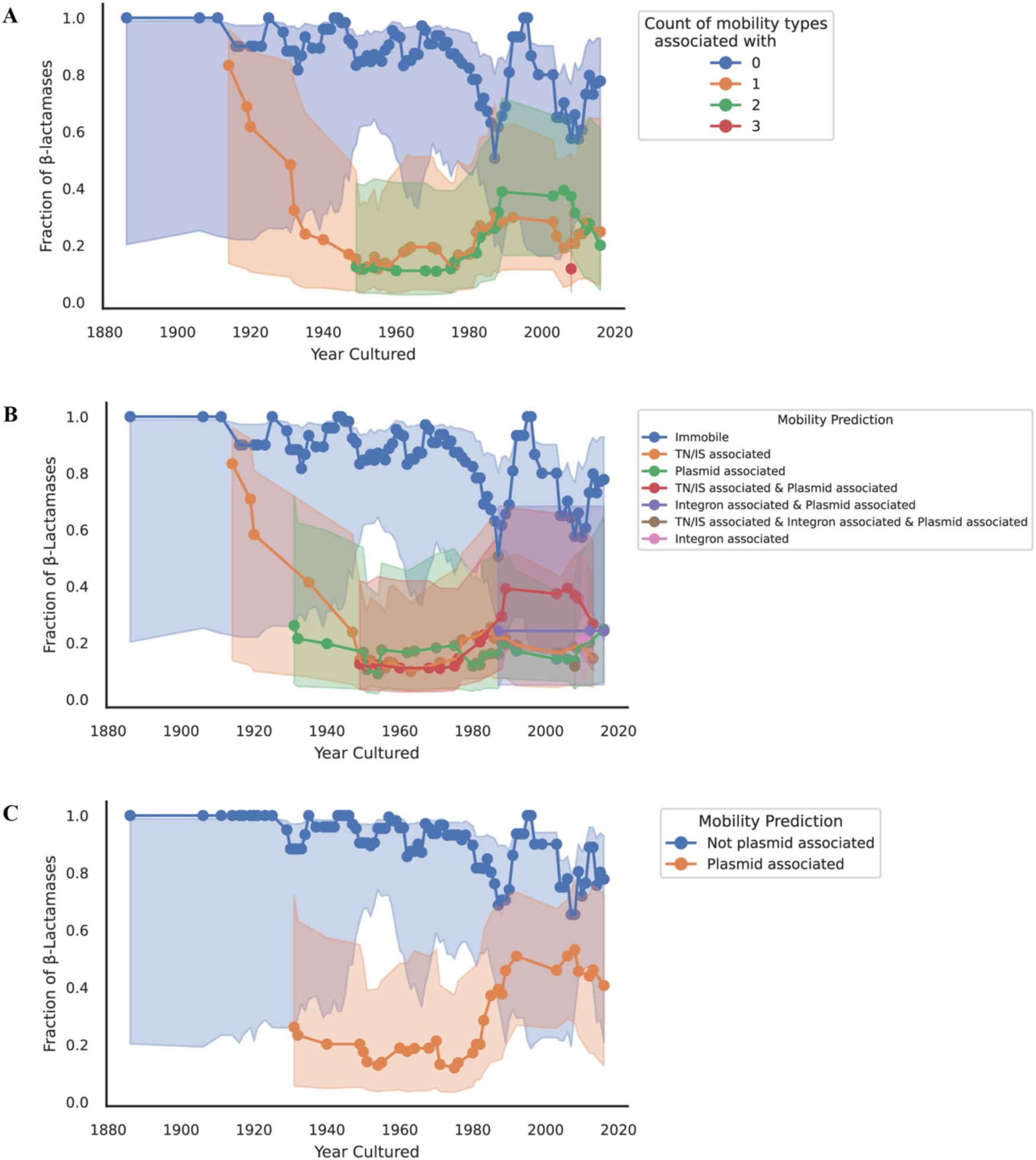
Beta-lactamases experience increasing mobility over the years, with most mobility driven by plasmids. Fraction of beta-lactamases classified as mobile over the course of the NCTC, shaded regions represent 95% confidence intervals based on sampling size (A) Number of mobility types associated with each beta-lactamase in the NCTC. (B) Breakdown of which combination of mobility types cause the counts in (A). (C) Fraction of beta-lactamases with some or no plasmid association.

**Supplementary Figure 7:**
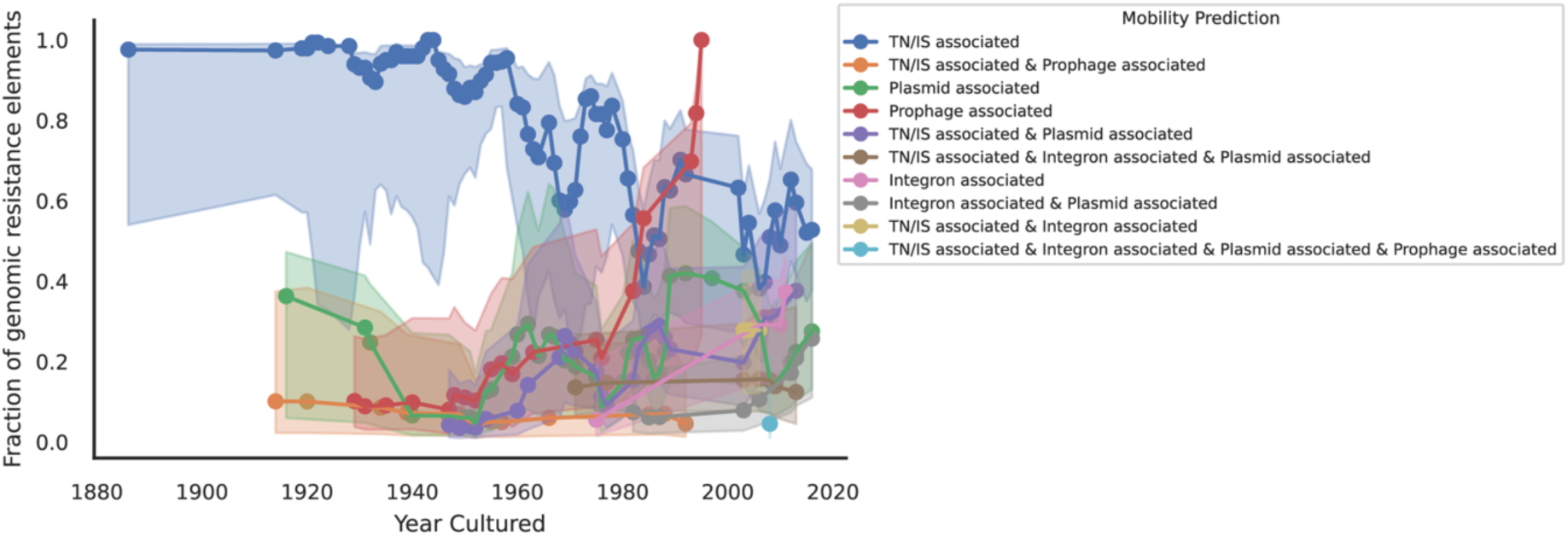
Increase in mobility access of mobile resistance elements appears to be driven by an increase in Plasmid-associated resistance elements. Fraction of mobile resistance elements as a function of time and the specific combination of mobility types observed. The shaded regions denote 95% confidence intervals from a beta distribution.

